# *Enterococcus faecalis* redox metabolism activates the unfolded protein response to impair wound healing

**DOI:** 10.1101/2025.08.07.669218

**Authors:** Aaron Ming Zhi Tan, Cenk Celik, Stella Yue Ting Lee, Mark Veleba, Caroline S. Manzano, Rahim MK Abdul, Guillaume Thibault, Kimberly A. Kline

## Abstract

*Enterococcus faecalis* is an opportunistic pathogen that thrives in biofilm-associated infections and delays wound healing, yet how it impairs host tissue responses is unclear. Here, we identified extracellular electron transport (EET) as a previously unrecognized source of ROS in *E. faecalis* and show that this activity directly triggers the unfolded protein response (UPR) in epithelial cells and delays epithelial cell migration. ROS detoxification with catalase suppressed *E. faecalis*-induced UPR and rescued epithelial cell migration, while exogenous H_2_O_2_ was sufficient to restore UPR activation in EET-deficient strains. Importantly, UPR disruption by pharmacological inhibition also impaired cell migration, highlighting a critical role for UPR homeostasis in wound repair. Our findings establish EET as a novel virulence mechanism that links bacterial redox metabolism to host cell stress and impaired repair, offering new avenues for therapeutic intervention in chronic infections.

## INTRODUCTION

*Enterococcus faecalis* is a gut commensal and opportunistic pathogen that causes difficult-to-treat biofilm-associated infections, including catheter-associated urinary tract infection, infective endocarditis, and chronic wound infections^1,2^. In wound settings, *E. faecalis* infection is associated with delayed epithelial migration and immune dysregulation^3^. The success of *E. faecalis* in these environments is often attributed to its metabolic adaptability, including survival under nutrient limitation and oxidative stress^4^. However, the extent to which *E. faecalis* metabolism actively interferes with host repair mechanisms is poorly understood.

One way that host cells respond to environmental and infection-induced insults is through the unfolded protein response (UPR), an evolutionarily conserved signalling pathway triggered by endoplasmic reticulum (ER) stress. The UPR integrates signals related to protein misfolding, membrane perturbations, and redox imbalance to restore homeostasis or induce apoptosis if stress persists^5–8^. Pathogens have evolved diverse strategies to manipulate the host UPR, often targeting its three main pathways (IRE1, PERK, and ATF6) to subvert host cell function, modulate immune responses, or even exploit UPR-regulated products as a nutrient source^9–14^. While some bacterial toxins and effectors can induce the UPR, in most cases, the microbial mechanisms by which the UPR is activated or dysregulated are undefined^11,15–17^.

*E. faecalis* generates substantial extracellular reactive oxygen species (ROS), including superoxide and hydrogen peroxide, in the absence of aerobic respiration or fumarate reduction^18,19^. These ROS can inflict DNA damage and tissue injury in infection models, as well as modulate host signalling pathways including redox-sensitive pathways like the UPR^20–22^. These ROS can inflict DNA damage and tissue injury in infection models, as well as modulate host signalling pathways including redox-sensitive pathways like the UPR^20,21,23^. However, the bacterial source of ROS in *E. faecalis* and its mechanistic consequences for host cell function have not been fully elucidated.

In this study, we show that *E. faecalis* activates the host UPR during wound infection. Through a forward genetic screen and functional validation, we identify extracellular electron transport (EET) as a previously unrecognised mechanism by which *E. faecalis* generates ROS, which in turn activates the UPR in epithelial cells and impedes their migration following wounding. We show that EET and associated demethylmenaquinone (DMK) biosynthesis pathways are required for superoxide and hydrogen peroxide generation, mutants of which produce less ROS, fail to activate the UPR, and do not impair epithelial cell migration. These findings not only establish a novel function for EET in ROS generation but also through its interaction with host UPR, as a novel metabolic virulence mechanism by which *E. faecalis* disrupts epithelial repair, thereby presenting new opportunities for targeting chronic *E. faecalis*-driven pathologies.

## RESULTS

### *E. faecalis* infection activates the UPR in a mouse model

We previously showed that *E. faecalis* infection impairs wound healing^3^. To gain insight into the mechanisms that influence delayed wound repair, we re-analysed our published single-cell RNA-seq dataset of *E. faecalis* infected mice wounds at 4 days post-infection^22^ (dpi) (**Figure 1A**, GSE229257). For each class we calculated enrichment scores for a panel of stress-response signatures^24^ (**Table S1**). Gene-set enrichment for canonical UPR targets revealed that the ER stress-response is not global but concentrated in immune cells (macrophages and neutrophils) and, most strikingly, in an infection-specific cluster of keratinocytes (**Figure 1B and 1C**). By contrast, oxidative-stress response (OSR) genes were upregulated not only in UPR-elevated keratinocytes and immune cells but also in fibroblasts (**Figure 1D, S1A, and S1B**), consistent with high fibroblast redox activity in infected tissue. We previously found that *E. faecalis* infection interferes with wound closure signatures, drives a partial epithelial-to-mesenchymal transition (EMT) in keratinocytes, and skews macrophages toward an anti-inflammatory phenotype^22^. The robust UPR response in keratinocytes offers a mechanistic clue where excessive ER stress could amplify the infection-induced EMT shift in these cells, undermining their migratory role and thereby hindering wound repair during *E. faecalis* infection.

**Figure 1.**
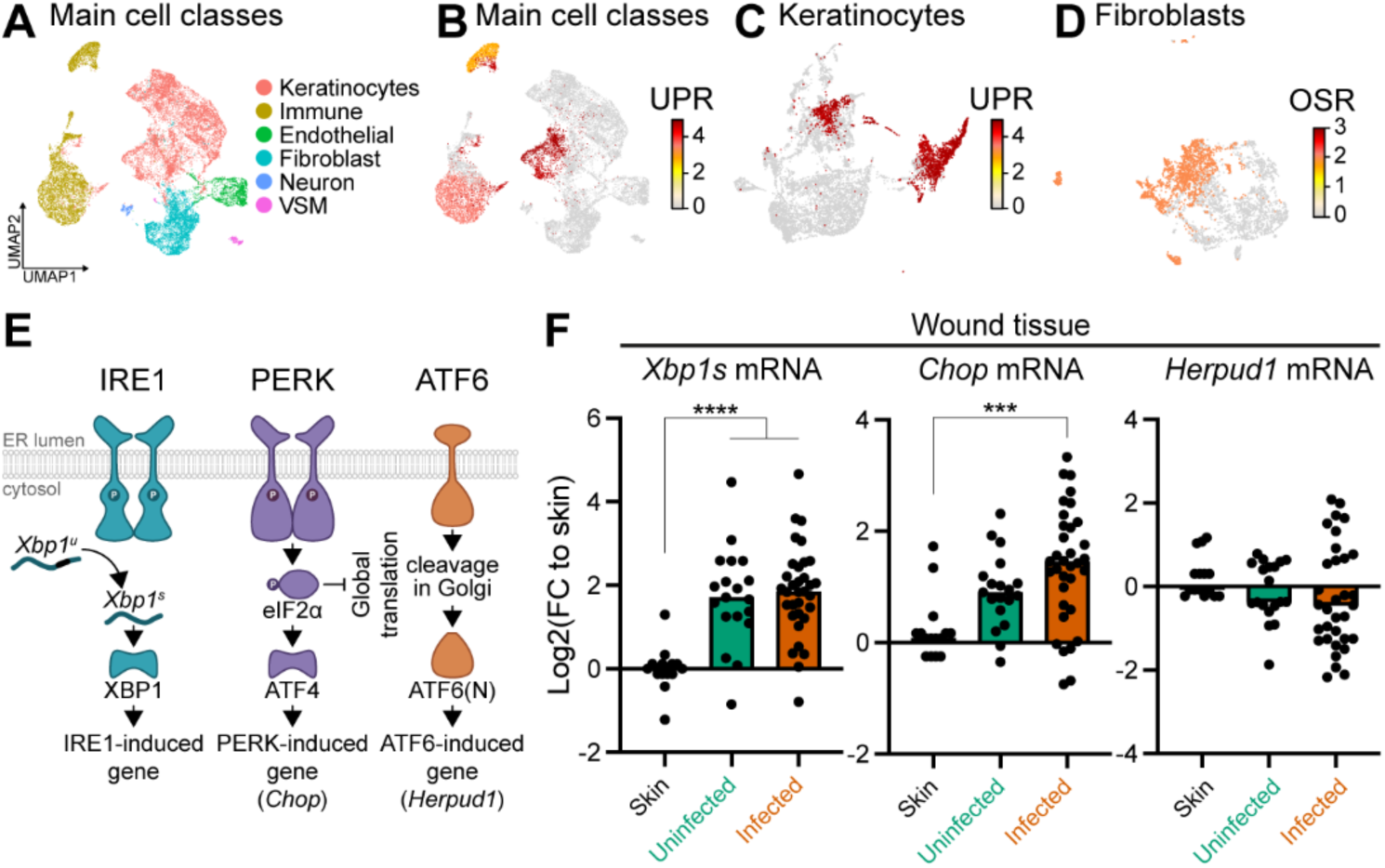
*E. faecalis* infection activates the UPR in a mouse model. **(A)** Uniform manifold approximation and projection (UMAP) of ∼23,000 single-cell transcriptomes from uninfected and *E. faecalis*-infected wounds^22^, recoloured here into the six broad cell classes used for downstream stress-response analyses. **(B)** Per-cell enrichment score for a curated unfolded protein response (UPR) gene set projected onto the UMAP in (A). **(C)** Same UPR enrichment as in (B) but displayed only for keratinocyte clusters. **(D)** Enrichment of an oxidative stress response (OSR) gene set plotted for fibroblast clusters. **(E)** Schematic representation of the UPR in mice, where spliced *Xbp1* (*Xbp1s*), *Chop*, and *Herpud1* serve as downstream markers of the IRE1, PERK, and ATF6 pathways, respectively. **(F)** Gene expression of UPR markers (*Xbp1s*, *Chop*, and *Herpud1*) at 6 dpi for uninfected and *E. faecalis*-infected 6–7 week-old C57BL/6J mouse skin wounds, normalized to intact skin. Significance was determined using one-way ANOVA, Dunnett’s test (unwounded skin, *n* = 16; wounded, uninfected, *n* = 20; wounded, infected *n* = 33; \*\*\**p* < 0 .001, \*\*\*\**p* < 0.0001).

To corroborate the scRNA-seq findings, we infected full thickness excisional wounds in mice with *E. faecalis* strain OG1RF and enumerated bacterial colony forming units (CFU) from the wounds at 6 days post-infection (6 dpi), chosen to target the proliferation and remodelling phase of healing, where we observed a 2-log reduction in *E. faecalis* bacteria burden in wounds compared to the starting inoculum, with no significant animal weight loss or attrition, consistent with our previous studies^3^ (**Figure S1C and S1D**). We quantified UPR-associated mRNA levels of spliced *Xbp1* (*Xbp1s*), *Chop*, and *Herpud1* as markers of Ire1, Perk1, and Atf6 activity, respectively, in whole wound tissue (**Figure 1E**). *Xbp1s* transcripts were significantly higher in uninfected wounds at 6 dpi compared to unwounded skin, indicating that the UPR is activated in wounds regardless of infection state (**Figure 1F**). A similar observation was reported in wounded transgenic mice expressing a XBP1-Luc fluorescence marker 8-10 days post-wounding^25^. By contrast, both *Xbp1s* and *Chop* transcript levels were significantly higher in *E. faecalis*-infected wounds at 6 dpi compared to unwounded skin samples, while *Herpud1* levels remained unchanged, indicating a lack of ATF6 activation (**Figure 1F**). Taken together, these data suggest that *E. faecalis* infection results in UPR dysregulation, which could impact normal wound healing^26^.

### IRE1 activation by *E. faecalis* impedes keratinocyte migration *in vitro*

To corroborate our *in vivo* and *in silico* findings, we examined UPR activation in keratinocytes (HaCaT) and fibroblasts (NIH-3T3) which are the dominant cell types in healthy skin that contribute to wound healing^27^*. E. faecalis* infection significantly increased mRNA expression of all three UPR pathway markers in NIH-3T3 cells (**Figure 2A**), whereas in HaCaT cells only *XBP1s* and *CHOP* were significantly upregulated (**Figure 2B**). As a positive control, we treated both cell lines with tunicamycin (Tm), which induces the UPR by inhibiting protein glycosylation in the ER leading to an accumulation of unfolded proteins^28^. Tm treatment significantly upregulated all three UPR pathway markers in both cell lines to a greater extent compared to *E. faecalis* infection. Since IRE1 is the most evolutionarily conserved branch of the UPR, we further examined its activation by *E. faecalis* by assessing the expression of *XBP1s* target genes^29,30^. These included the ER chaperone BiP (encoded by *HSPA5*) and EDEM1, which promote ER homeostasis and are hallmarks of IRE1 activation^31,32^. Infected cells exhibited a significant increase in *EDEM1* transcripts along with elevated levels of XBP1s and BiP proteins (**Figure 2C and 2D**), confirming that *E. faecalis* activates the conserved IRE1 pathway *in vivo* and *in vitro*.

**Figure 2.**
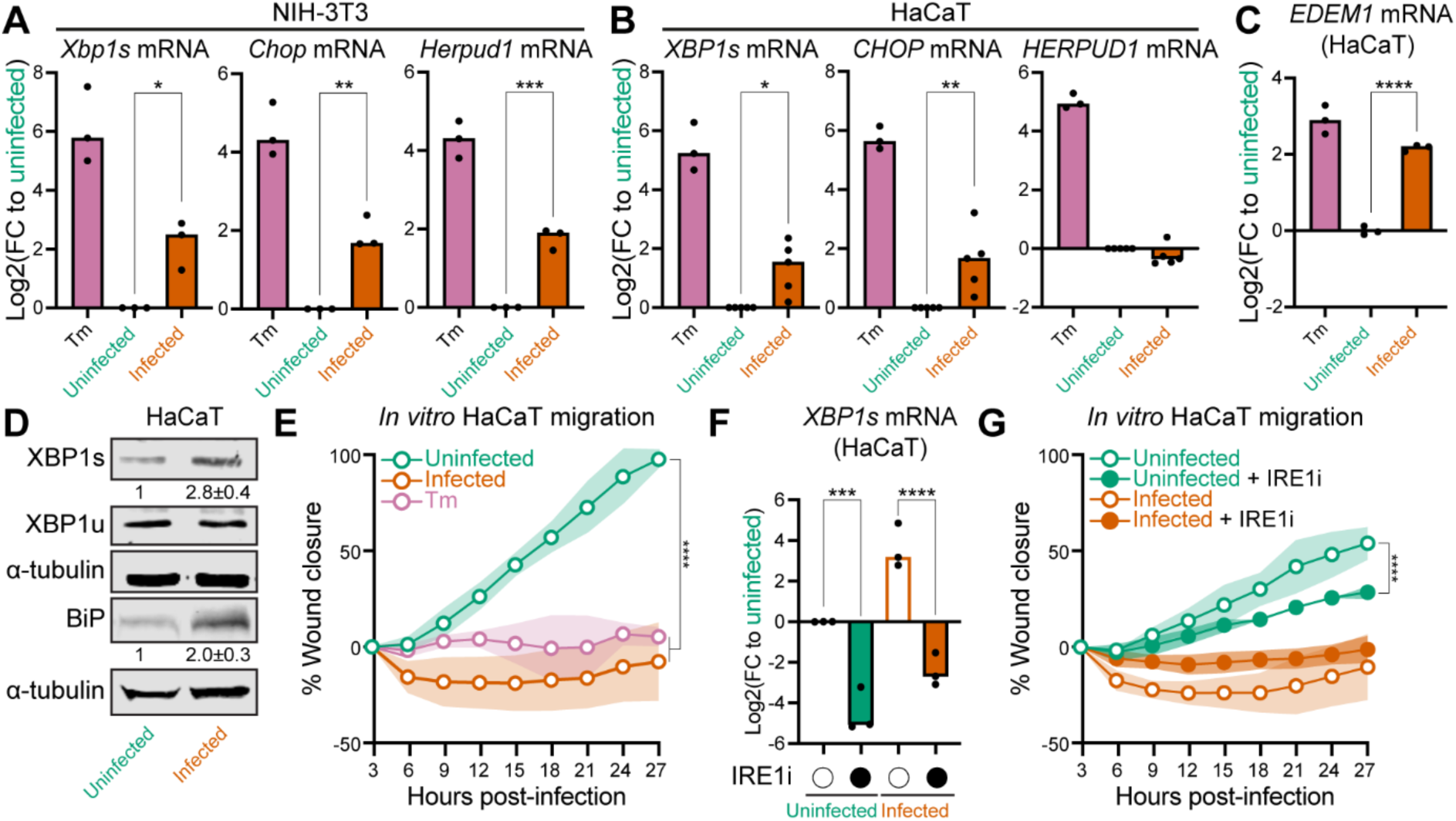
IRE1 activation by *E. faecalis* impedes keratinocyte migration *in vitro*. **(A-B)** Gene expression of UPR markers (*Xbp1s, Chop, and Herpud1*) in infected with *E. faecalis* at MOI of 800 (A) or 600 (B) or tunicamycin (Tm) treated NIH-3T3 mouse fibroblasts (A) or HaCaT human keratinocytes (B) (*n* = 3). See Methods for MOI optimization description. **(C)** Gene expression of IRE1 downstream gene (*EDEM1*) in cells treated as in (A) (*n* = 3). **(D)** Quantitative analysis and representative immunoblots showing levels of XBP1s and BiP in HaCaT cells treated as in (A). **(E)** Scratch wound assay quantification for uninfected, infected and Tm-treated cells (positive control) (*n* = 4 biological replicates). **(F)** Gene expression of *XBP1s* from HaCaT cells treated with the DMSO control (open circles) or the IRE1 inhibitor (IRE1i) 4µ8c (closed circles) (*n* = 3, one-way ANOVA, Dunnett’s test) **(G)** Scratch wound assay quantification for uninfected and infected cells treated with 0.5% DMSO control (open circles) or IRE1i (close circles) (*n* = 4 biological replicates). Significance was determined using one-way ANOVA Dunnett’s test (A, B, and F) or two-way ANOVA Tukey’s test (E and G) (\**p* < 0.05, \*\**p* < 0.01, \*\*\**p* < 0 .001, \*\*\*\**p* < 0.0001).

We next investigated the impact of infection-induced UPR on wound closure using an *in vitro* HaCaT scratch wound assay. By 15 hpi, both *E. faecalis*-infected and Tm-treated cells exhibited significantly slower migration compared to uninfected cells, with the difference becoming more pronounced by 27 hpi (**Figure 2E and S2A; Video S1**). Notably, neither infected nor Tm-treated cells displayed appreciable migration throughout the assay period, and infected cells even showed signs of wound edge retraction as early as 6 hpi, consistent with cell shrinkage or detachment which can be early indicators of apoptosis. To determine whether UPR induction via IRE1 was responsible for impaired migration, we treated both uninfected and infected cells with IRE1 inhibitor (4µ8c), which blocks the RNase activity of IRE1^33^ (IRE1i). Treatment with 50 µM IRE1i was sufficient to block *E. faecalis*-induced IRE1 in HaCaT cells (**Figure 2F**). IRE1i also slowed migration in uninfected wells, with the delay evident by 18 hpi and more pronounced at 27 hpi. (**Figure 2G and S2B; Video S2**). In infected cells, migration was minimal regardless of IRE1i treatment, showing no significant migration at 27 hpi relative to the 3 hpi baseline.

We did not use proliferation inhibitors, such as mitomycin C, which are typically used to differentiate between proliferation and migration in these assays, because the compound caused widespread cell detachment in infected cells during preliminary experiments. Nonetheless, we attribute the observed wound closure primarily to cell migration rather than proliferation for several reasons: (i) the short 24-h duration of the assay, (ii) the long doubling time of confluent HaCaT cells (approx. 32-36 h), and (iii) previous studies have established that migration is the dominant factor in similar short-term scratch assays^26^. Thus, our findings suggest that neither UPR hyperactivation (during infection) nor hypoactivation (after IRE1 inhibition) alone fully explains keratinocyte migration; rather, both extremes of UPR activity are implicated.

### Extracellular electron transport drives *E. faecalis* UPR activation and migration arrest

To dissect how *E. faecalis* induces the UPR, we designed a high-throughput assay for UPR induction: a NIH-3T3 reporter line (3T3R) that fluoresces when IRE1 splices a 26-nt intron from a truncated human *XBP1* fused to *mApple* (**Figure 3A and 3B**). While IRE1 is quiescent, a premature stop codon between *XBP1* and *mApple* blocks translation, causing cells to remain non-fluorescent. When IRE1 is activated, intron excision shifts the reading frame, removes the stop codon, and allows production of the full XBP1s-mApple fusion, generating a red signal quantifiable by wide-field microscopy. Using this reporter, we screened a defined transposon (Tn) library of 14,976 *E. faecalis* OG1RF mutants (**Figure 3A**) to identify mutants that failed to trigger UPR activation.

**Figure 3.**
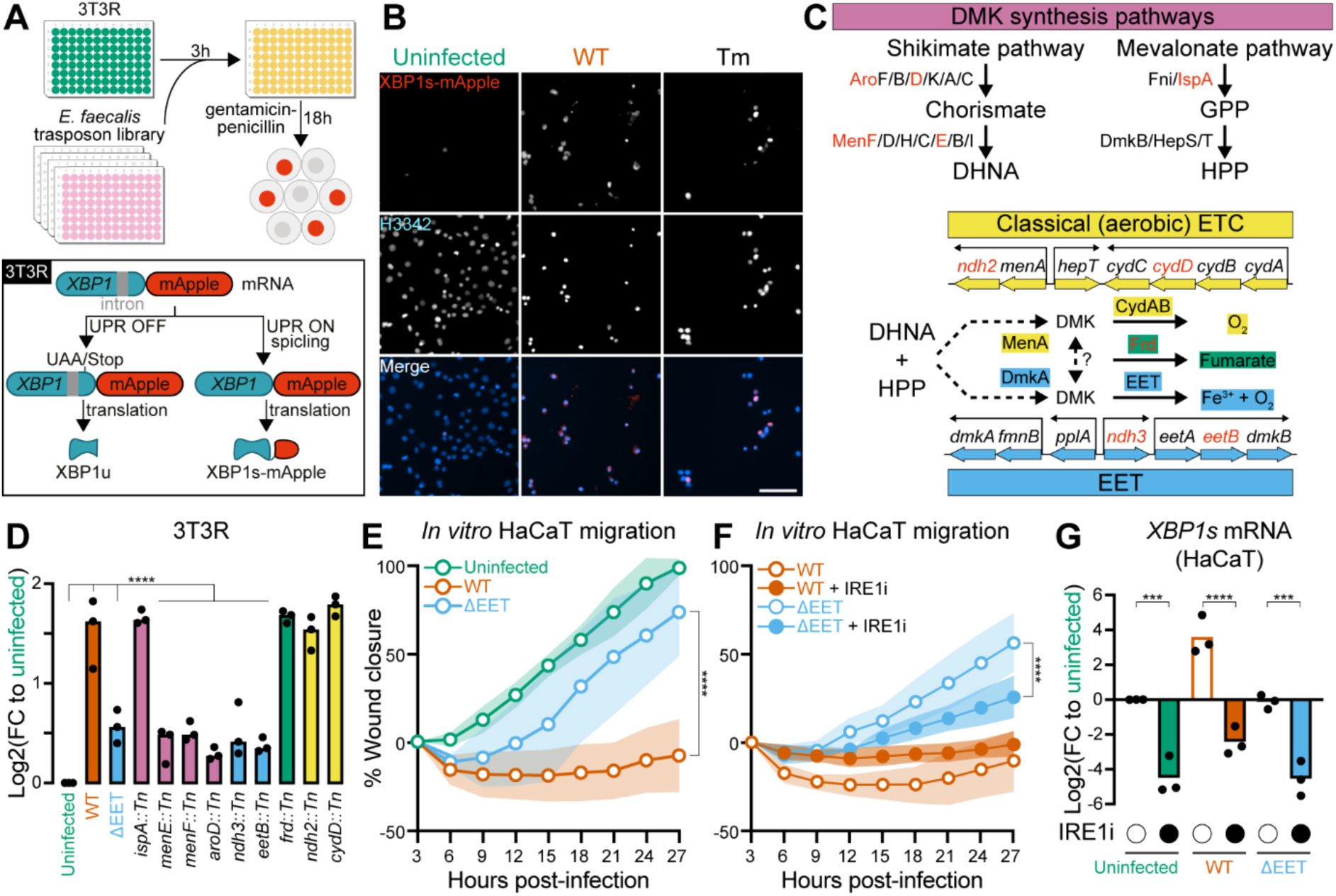
Extracellular electron transport drives *E. faecalis* UPR activation and migration arrest. **(A)** *E. faecalis* OG1RF transposon screen in which a library of 14976 mutants was screened against a NIH-3T3 cell line expressing the Xbp1-mApple reporter system (3T3R). **(B)** Representative epifluorescnece microscopy images of 3T3R under different conditions (uninfected, WT, and Tm-treated) at 21 hpi. Scale bar, 100 μm. **(C)** Diagram showing the pathways in which a subset of UPR defective mutants (genes/proteins with red font) were identified. **(D)** Validation of UPR defective mutants with 3T3R (*n* = 3). **(E-F)** Scratch wound assay quantification for HaCaT infected with (E) WT and ΔEET, which were also (F) treated with either 0.5% DMSO control (open circles) or IRE1i (close circles). Uninfected and WT data are identical to Figure 2E and 2F and replicated here for ease of comparison (*n* = 4 biological replicates). **(G)** UPR induction at 24 hpi in HaCaT after treatment with 0.5% DMSO control (open circles) or IRE1i (closed circles) under uninfected, WT-infected, and ΔEET-infected conditions. Uninfected and WT-infected findings are identical to Figure 2F and replicated here for ease of comparison (*n* = 3). Significance was determined using one-way ANOVA Dunnett’s test (D and G) or two-way ANOVA Tukey’s test (E and F) (\*\*\**p* < 0 .001, \*\*\*\**p* < 0.0001).

Applying an upper threshold of 30% XBP1s-mApple positive (XBP1s+) cells in a given population, we identified 457 UPR defective mutants, corresponding to 369 unique genetic loci (**Figure S3A**). We filtered out mutants with multiple transposon insertions, insertions outside of coding regions, and those with significant growth defects after overnight culture in BHI media. Pathway analysis did not show consistent enrichment patterns; however, we noticed that a substantial number of mutants had insertions in genes associated with carbohydrate catabolism and respiration, key components of redox metabolism. This observation prompted us to focus on pathways involved in the synthesis of components and co-factors that facilitate electron flow from carbohydrate catabolism to terminal electron acceptors. Transposon insertions mapped to three respiratory pathways of *E. faecalis*: aerobic respiration (*ndh2, cydD*), fumarate reduction (*frd*), and extracellular electron transport (EET) (*ndh3*, *eetB*) (**Figure 3C**). Additional insertions were identified in genes involved in the biosynthesis of the quinone electron carrier demethylmenaquinone (DMK) (*menF, menE*), as well as upstream precursors such as chorismate and geranyl pyrophosphate, generated either by the shikimate pathway (*aroD*) or the mevalonate pathway (*ispA*). DMK is an integral part of all three respiratory pathways by mediating electron transfer between their membrane-associated components.

We validated mutants of these respiratory pathways identified in the primary screen and found that only mutants disrupted in quinone electron carrier synthesis (*menE, menF, aroD),* and EET (*ndh3, eetB*) displayed significantly reduced UPR induction compared to wild-type *E. faecalis* (WT) (**Figure 3D**). Since these genes are involved in central metabolic processes, we quantified the growth of each mutant to ensure that reduced UPR induction was not simply due to impaired bacterial replication during infection. However, all Tn mutants grew similarly to WT in cell culture media (**Figure S3B and S3C**), eliminating growth defects as a confounding factor. Given the strong association between EET and UPR induction, we also tested a deletion mutant lacking entire EET operon (ΔEET) (**Figure 3C and 3D**). As expected, the ΔEET mutant was defective in UPR induction, yet retained growth and antibiotic susceptibility profiles similar to the parental OG1RF WT strain (**Figure S3B-S3D)**. To rule out the possibility that lower UPR induction by ΔEET could be due to higher cytotoxicity causing cell loss resulting in weaker XBP1s-mApple fluorescent signals, we assessed cytotoxicity in HaCaT cells at 3 and 24 hpi following infection with WT and ΔEET. There was no difference in cytotoxicity between uninfected, WT- or ΔEET-infected cells at 3 hpi (**Figure S3E**). However, WT but not ΔEET-infected cells had significantly higher cytotoxicity compared to uninfected cells at 24 hpi. The significantly higher cytotoxicity in WT-infected cells at a delayed timepoint of 24 hpi but not immediately after infection at 3 hpi suggests that WT infection was not directly causing cell death but rather indirectly via UPR hyperactivation. The lack of difference at 24 hpi between uninfected and ΔEET-infected cells also confirms the UPR defective nature of ΔEET, as delayed cell death is a hallmark characteristic of chronic UPR hyperactivation^34,35^. To further confirm that the loss of UPR induction was specific to the EET pathway and not to secreted toxins, as is the case for Group A Streptococcus^14,36^, we tested a deletion mutant of the Type 7 secretion system (ΔT7SS), encoding predicted secreted toxins^37^. The ΔT7SS mutant induced the UPR to the same extent as WT (**Figure S3F**), supporting the conclusion that the phenotype of the ΔEET mutant is a direct consequence of its function in redox metabolism.

Based on these characteristics, ΔEET was selected as the model UPR defective mutant for downstream studies. We used this mutant to assess whether *E. faecalis* UPR induction via the EET pathway contributes to the inhibition of keratinocyte cell migration. Unlike WT infection, ΔEET did not significantly impair migration, instead showing similar migration, comparable to uninfected controls (**Figure 3E and S3G; Video S3**). However, treating ΔEET-infected cells with IRE1i resulted in significantly slower migration at the 27 hpi endpoint (**Figure 3F and S3H**). These findings suggest that lack of UPR induction by ΔEET allows for physiological levels of UPR induction that support normal cell migration. Furthermore, IRE1i treatment induces UPR hypoactivation in ΔEET-infected cells (**Figure 3F and S3H; Video S4**), dysregulating UPR homeostasis and impairing cell migration, similar to that observed in uninfected cells treated with IRE1i (**Figure 2G**, **Figure 3G**). The recovery of cell migration with ΔEET infection and its reversal by IRE1i treatment demonstrate that *E. faecalis* EET is associated with UPR induction and impaired cell migration.

### EET-derived ROS is sufficient to activate the UPR, disrupting epithelial migration

Disruption of DMK synthesis in *E. faecalis* impairs both EET function and extracellular superoxide generation^18,38^. OG1RF mutants disrupted in genes involved in quinone electron carrier synthesis (*aroE, aroC, aroA, menB, menD, menE*) produce less superoxide (O_2_^•–^) than WT^19^. Superoxide radicals undergo pH-dependent spontaneous dismutation to generate hydrogen peroxide (H_2_O_2_) which can then participate in Fenton chemistry to generate hydroxyl radicals and other ROS in the presence of transition metals (**Figure 4A**). H_2_O_2_ has been shown to induce the UPR in myotubule and epithelial cells at 50 µM and 200 µM, respectively^20,23^. However, this may not be a universal response among host cells as another group reported no significant increase in *XBP1s* expression for fibroblast when treated with 1 mM H_2_O_2_^39^. We hypothesised that superoxide generation via EET drives UPR induction in epithelial cells via H_2_O_2_. To test whether H_2_O_2_ alone is sufficient to induce the UPR in our models, we treated 3T3R cells with increasing concentrations of H_2_O_2_ for 3 hours followed by a 21-hour recovery period. XBP1s fluorescence increased significantly at 250 and 500 µM **(Figure 4B**). This finding confirms that NIH-3T3 cells can mount a UPR in response to H_2_O_2_, supporting the idea that ROS generated via EET contributes to UPR induction during infection.

**Figure 4.**
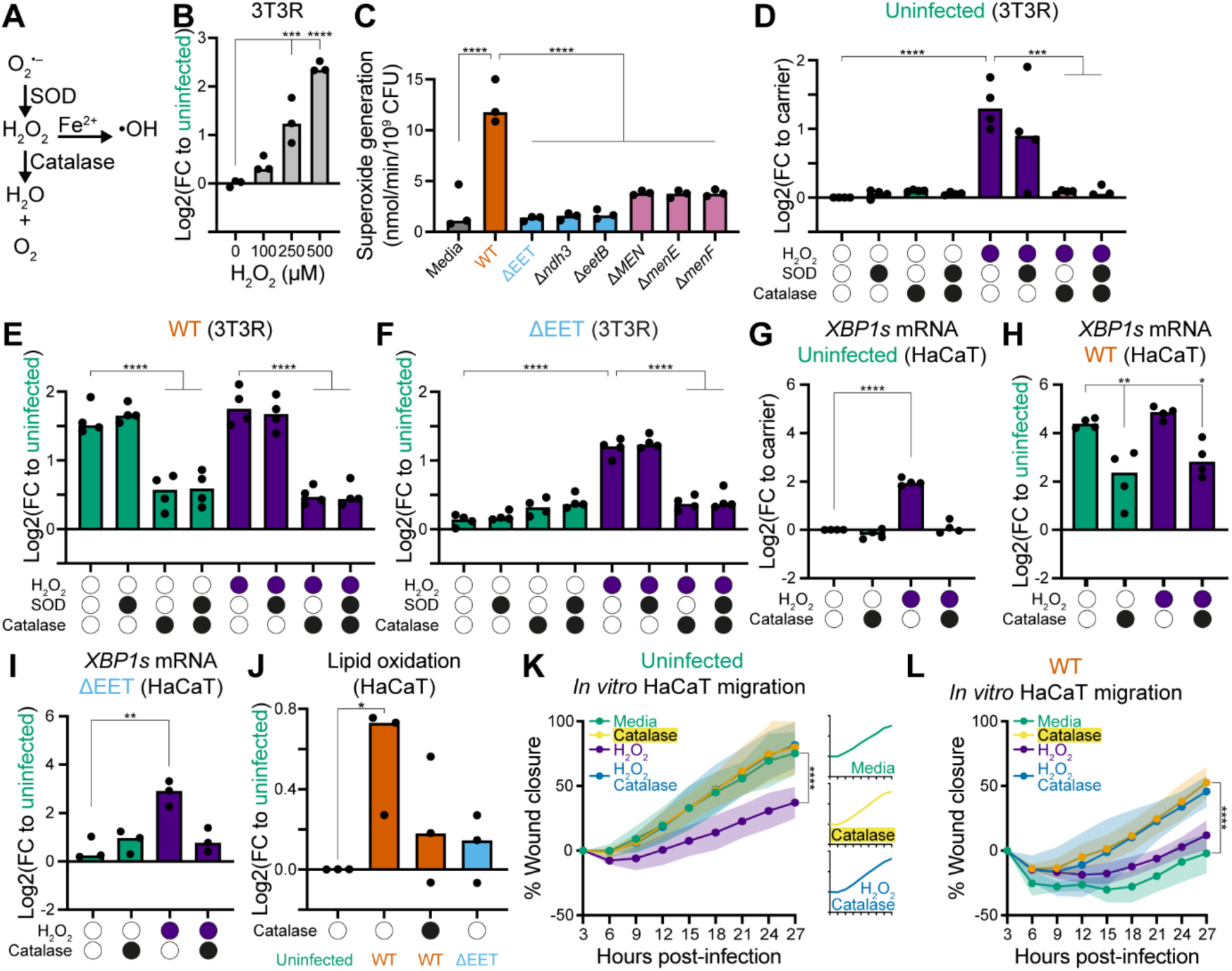
EET-derived ROS is sufficient to activate the UPR, disrupting epithelial migration. **(A)** Diagram showing the dismutation of superoxide radical (O_2_^•–^) into hydrogen peroxide (H_2_O_2_) via the catalytic activity of superoxide dismutase (SOD), or spontaneously via a pH-dependent process. H_2_O_2_ can be converted into a highly reactive hydroxyl radical in the presence of transition metals like ferrous ions (Fe^2+^) via the Fenton reaction or neutralised into water and oxygen if catalase is present. **(B)** Dose-dependent H_2_O_2_ mediated UPR induction in 3T3R (*n* = 3). **(C)** In-frame deletion mutants corresponding to the Tn mutants of the EET and DMK synthesis pathways which were identified as UPR defective hits during the transposon screen. The ΔMEN mutant had its entire operon deleted, which contained the *menF/D/C/E/B* genes (*n* = 3). **(D-F)** UPR induction in 3T3(R) for (D) uninfected (E) WT infection, and (F) ΔEET infection, in the presence (closed circles) or absence (open circles) of H_2_O_2_, SOD, and catalase (*n* = 4). **(G-I)** UPR induction in uninfected (G), WT infected (H), and ΔEET infected (I) HaCaT cells in the presence (closed circles) or absence (open circles) of H_2_O_2_ and catalase (*n* ≥ 3). **(J)** Lipid oxidation in uninfected, WT infected, and ΔEET infected HaCaT cells using BODIPY 581/591 C11 (*n* = 3). **(K-L)** Scratch wound assay quantification in uninfected (K) or WT infected (L) HaCaT cells treated with catalase and/or H_2_O_2_ (*n* = 3 biological replicates). Significance was determined using one-way ANOVA Dunnett’s test (B-J) or two-way ordinary ANOVA, Tukey’s multiple comparisons test (K-L) (\**p* < 0.05, \*\**p* < 0.01, \*\*\**p* < 0 .001, \*\*\*\**p* < 0.0001).

Next, to test whether EET indeed generates ROS, we quantified superoxide generation by validated UPR-defective mutants (ΔEET, and transposon insertion mutants in *menE, menF, arodD, ndh3, eetB*). At the same time, we generated in-frame deletion mutants for each of these genes. All exhibited significantly lower superoxide generation compared to WT and non-UPR defective mutants (*ispA, frd, ndh2, cydD*), and the in-frame deletion mutants were similar to their respective transposon mutants **(Figure 4C and S4A)**, supporting a link between EET-dependent superoxide production and UPR activation.

Superoxide dismutase (SOD) and catalase are antioxidants that catalyse the dismutation of superoxide radicals into H_2_O_2_ or H_2_O_2_ into water and oxygen, respectively (**Figure 4A**). To determine whether *E. faecalis-*derived ROS drives UPR induction, we treated infected 3T3R cells with exogenous SOD and catalase. In control experiments, neither enzyme altered XBP1s expression in uninfected cells, whereas treatment with 250 µM H_2_O_2_ to simulate ROS-induced stress robustly increased XBP1s expression (**Figure 4D**). In cells stimulated with H_2_O_2_, the addition of catalase significantly reduced XBP1s expression, whereas SOD had no effect. This result confirms that H_2_O_2_ is the specific reactive oxygen species driving UPR induction in this assay.

In WT-infected cells, catalase again significantly reduced XBP1s expression, while SOD had no effect (**Figure 4E**). Adding H_2_O_2_ to WT-infected cells did not further increase UPR activation, suggesting that *E. faecalis*-generated ROS levels are already saturating whereas co-treatment with catalase reversed this effect. By contrast, catalase had no impact on ΔEET-infected cells, consistent with their low ROS generation (**Figure 4F**). However, functional complementation of the ΔEET with H_2_O_2_ partially restored UPR induction, which was again reversed by catalase, confirming that H_2_O_2_ is sufficient to rescue the UPR defective phenotype when EET is disrupted. These findings were replicated in HaCaT cells (**Figure 4G-4I**). Since lipid peroxidation is a known cause of UPR activation^40^. We next measured it in infected cells using a BODIPY 581/591 C11 fluorescent probe. WT infected cells exhibited significantly more lipid peroxidation than uninfected controls, while cells infected with the ΔEET mutant showed no significant change (**Figure 4J**). These data support a model where EET-derived ROS from *E. faecalis* induces the UPR via lipid peroxidation.

To determine whether *E. faecalis* generated ROS impairs wound healing, we assessed *in vitro* wound closure after treatment with H_2_O_2_ or/and catalase. H_2_O_2_ treatment of uninfected cells significantly slowed cell migration at 27 hpi, although no wound edge retraction was observed, unlike in WT-infected cells (**Figure 4K and S4B; Video S5**). Co-treatment with catalase restored cell migration to physiological levels. Similarly, catalase treatment of WT-infected cells significantly improved cell migration, which approached that of uninfected controls and without retraction at 27 hpi (**Figure 4L and S4C; Video S6**). Adding H_2_O_2_ to untreated WT-infected cells had no further effect, consistent with UPR saturation seen earlier (**Figure 4E and 4H**). Finally, co-treatment with H_2_O_2_ and catalase mirrored catalase-treatment alone, reinforcing the role of *E. faecalis*-derived H_2_O_2_ in cell migration arrest (**Figure 4L and S4C; Video S6**). Altogether, these findings demonstrate that the EET pathway of *E. faecalis* generates ROS which oxidises lipids in epithelial cells, hyperactivating the UPR, and inhibiting cell migration.

## DISCUSSION

Our findings identify *E. faecalis* extracellular electron transport (EET) as a novel virulence mechanism that links bacterial redox metabolism to host stress responses and impaired tissue repair. We show that EET-dependent production of reactive oxygen species (ROS), particularly H_2_O_2_, activates the unfolded protein response (UPR) in epithelial cells. UPR dysregulation by *E. faecalis* disrupts normal epithelial function and cell migration, revealing a direct mechanistic connection between bacterial energy metabolism and host healing processes.

*E. faecalis* is known to produce ROS, which have been previously linked to host cellular injury and even carcinogenesis^21^. In this study, we refine the genetic basis of this activity in terms of a dedicated electron transport system. Previous studies showed that *E. faecalis* generates extracellular superoxide through a process that requires demethylmenaquinone (DMK), which is diminished when terminal quinol/cytochrome oxidases are functional, suggesting that respiratory disruption changes the dynamics of electron flow to one that favours the reduction of oxygen to superoxide, resulting in ROS production^19^. In this model, NADH incompletely reduces DMK, producing semiquinone intermediates which, in the absence of classical aerobic or anaerobic respiration, donate electrons univalently to molecular oxygen, generating superoxide which spontaneously dismutate into hydrogen peroxide which can contribute to host oxidative stress^19^. This movement of electrons may be mediated by soluble shuttles, such as flavins^41,42^. In this study, we refine the mechanistic basis of *E. faecalis* ROS generation by identifying a critical role for extracellular electron transport (EET) in this process.

While disrupting aerobic respiration is required for ROS generation and EET function^19,38^, our data show that the disruption of aerobic respiration alone is not sufficient and that ROS generation also depends on an intact EET system. However, the mechanistic contribution of specific EET components involved in ROS generation remains to be determined. Whether these components overlap with those required for extracellular metal reduction or electrode respiration or represent a distinct branch of EET machinery activated under redox stress, remains an important question for future work.

The requirement for EET in *E. faecalis* ROS production aligns with growing evidence that extracellular electron transfer in Gram-positive bacteria extends beyond classical anaerobic respiration. While canonical diderm systems in general like Shewanella and Geobacter use cytochromes and conductive pili to transfer electrons to external acceptors^43,44^, recent studies show that monoderm bacteria, including *Listeria monocytogenes* and *Lactobacillus plantarum*, use flavin-based EET to support energy conservation, redox homeostasis, and virulence under host-relevant conditions^41,42,45^. In particular, *E. faecalis* and *L. monocytogenes* rely on EET for fitness in the mouse gastrointestinal tract^41,46^, and *L. plantarum* employs EET to enhance ATP yield via substrate-level phosphorylation in the absence of a classical respiratory chain^47^. Our findings position *E. faecalis* within this emerging framework and reveal a distinct facet: in the presence of oxygen within a heme-free environment, EET contributes to univalent electron transfer from reduced DMK to oxygen, generating superoxide and hydrogen peroxide. Unlike environmental microbial EET systems that avoid oxygen to prevent radical formation, *E. faecalis* appears to exploit this interaction, linking EET to oxidative stress at the host-pathogen interface.

The UPR is increasingly recognized as a central node in host stress responses during infection, particularly in epithelial and immune cells. Multiple pathogens including *Salmonella enterica*, *Helicobacter pylori*, *Pseudomonas aeruginosa*, *Streptococcus pyogenes,* and *Brucella melitensis* have been shown to manipulate the UPR through secreted toxins or effector proteins, often to promote intracellular survival or dampen immune responses^9–11,14,48^. Here we demonstrate that *E. faecalis* selectively activates the IRE1 and PERK arms of the UPR both *in vivo* and *in vitro*, manifesting in impaired epithelial cell migration. Unlike previously described examples, *E. faecalis* induces the UPR independent of dedicated virulence factors, instead leveraging metabolic ROS production via EET. Similar redox-based virulence strategies have been described in other pathogens. *S. pneumoniae* produces hydrogen peroxide via SpxB, contributing to epithelial damage^49,50^*. P. aeruginosa* phenazines generate intracellular ROS in airway cells^51,52^. Yet *E. faecalis* is the first example, to our knowledge, where a defined EET system is shown to drive ROS production that directly alters host stress signalling and function.

A compelling hypothesis arising from our data is that the observed UPR hyperactivation and impaired cell migration are consequences of ferroptosis. Ferroptosis is a regulated, iron-dependent form of cell death distinct from apoptosis, driven by the catastrophic accumulation of lipid peroxides. This process is typically restrained by the antioxidant enzyme GPX4, and its failure leads to membrane damage^53,54^. Ferroptosis is increasingly recognized as a critical factor in diverse pathologies and host-pathogen interactions^55–57^. Importantly, the accumulation of lipid peroxides, the biochemical hallmark of ferroptosis is a known trigger of severe endoplasmic reticulum stress, providing a direct mechanistic link to UPR activation^40,58,59^. Our findings align remarkably well with this framework. The lipid peroxidation we observed in infected keratinocytes is the defining feature of ferroptosis, and the subsequent cell retraction is a classic morphological correlate of this death pathway^53,54,60^. This model mechanistically connects *E. faecalis* EET-driven ROS production to the downstream cellular pathologies of lipid stress, UPR activation, and inhibition of cell migration. While our experiments with catalase confirm ROS as the primary trigger, they do not exclude ferroptosis as the ultimate executioner pathway. We therefore propose that the *E. faecalis*-host interaction may represent a novel model of infection-induced ferroptosis. Testing this hypothesis, for which our study provides the foundational rationale, will be a critical next step and could be directly addressed by employing specific inhibitors like ferrostatin-1.

Our data further show that UPR activation must be tightly regulated for effective wound healing. These findings are consistent with previous studies in aged keratinocytes and fibroblasts displaying higher baseline levels of UPR markers and slower *in vitro* wound cell migration, which could be reversed upon treatment with 4-phenylbutyrate, a broad-acting UPR inhibitor^26^. Similarly, we show that pharmacologic inhibition of IRE1, the key UPR sensor, led to hypoactivation and impaired cell migration even in uninfected cells, underscoring the importance of physiological UPR signalling during repair. Regulated UPR induction is also necessary for other wound healing processes like the differentiation of dermal fibroblasts to myofibroblasts which promote wound contracture and collagen deposition^61^. Furthermore, UPR inhibitors especially PERK inhibitors have demonstrated significant cytotoxicity to pancreatic islet cells which depend on mild UPR induction to perform their secretory function^62^. Even if UPR induction is not lowered to below baseline levels, UPR inhibition will still be counterproductive in restoring homeostasis in pathologies where host cells are already dependent on some level of UPR Induction to perform physiological processes. Therefore, targeting the source of UPR dysregulation i.e. bacterial ROS production, rather than host UPR, may constitute a more effective therapeutic strategy.

In summary, this work reveals that *E. faecalis* leverages its respiratory machinery not only for metabolic flexibility but also to perturb host cell physiology. While EET has been linked to efficient infection of the gastrointestinal tract, this work presents molecular details that may contribute to its role in pathogenesis. Future studies should examine the role of EET *in vivo*, its regulation and contribution within polymicrobial settings, and the potential for targeting redox metabolism to mitigate *E. faecalis* infections that are increasingly recalcitrant to antibiotic therapy.

## RESOURCES AVAILABILITY

### Lead contact

Further information and requests for resources and reagents should be directed to and will be fulfilled by the lead contact, Kimberly Kline.

### Materials availability

This study did not generate new unique reagents.

### Data and code availability

This paper did not generate any new original datasets nor codes.

Any additional information required to reanalyse the data reported in this paper is available from the lead contact upon request.

## ACKNOWLEDGEMENTS

We are grateful to Thibault and Kline lab members for helpful discussions and critical reading of the manuscript. We thank Gary Dunny and Jennifer Dale for providing the *E. faecalis* transposon mutants. We would also like to thank the Centre for Biomedical Informatics (Drs. James A. Miller and Bernett Teck Kwong Lee) and the NTU Optical Bio-imaging Centre (NOBIC) for their support. This work was supported by funds from the National Medical Research Council Open Fund (MOH-000566 to GT and KAK), the Singapore Ministry of Education Academic Research Fund Tier 1 (RG31/24 to GT), NTU Research Scholarship to SYTL and RMKA (predoctoral fellowship), and the Swiss National Science Foundation (SNSF grant 310030_212262 to KAK). Parts of this work were also supported by the National Research Foundation and Ministry of Education Singapore under its Research Centre of Excellence Program (SCELSE).

## AUTHOR CONTRIBUTIONS

Conceptualization, A.M.Z.T., G.T. and K.A.K.; Resources A.M.Z.T., M.V., C.S.M., and R.M.K.A.; Data curation, C.C. and R.M.K.A.; Formal analysis, A.M.Z.T., C.C., S.Y.T.L., and R.M.K.A.; Validation, A.M.Z.T.; Investigation, A.M.Z.T., C.C., S.Y.T.L.; Visualization, A.M.Z.T., C.C., S.Y.T.L. and G.T.; Methodology, A.M.Z.T., C.C., S.Y.T.L., G.T. and K.A.K.; Writing – original draft, A.M.Z.T., G.T. and K.A.K.; Writing – review and editing, A.M.Z.T., C.C., G.T. and K.A.K.; Project administration, G.T. and K.A.K.

## MATERIALS AND METHODS

### Bacterial strains and growth conditions

All bacterial strains used in this study are listed in **Table S2**. *E. faecalis* strains were routinely cultured on brain-heart infusion (BHI; Neogen #NCM0016A) agar plates and grown in BHI broth. *E. coli* strains, used for DNA isolation and plasmid manipulation, were cultured in Luria-Bertani (LB) broth or on LB agar plates at 37°C. When required, antibiotics were added at the following final concentrations: erythromycin (Em), 500 µg/ml for *E. coli* and 25 µg/ml for *E. faecalis*; rifampin (Rif), 25 µg/ml; and chloramphenicol (Cm), 10 µg/ml. To prepare overnight cultures, a single *E. faecalis* colony was inoculated into a 14 ml tube containing 4 ml of BHI broth, with the lid tightly sealed. Cultures were grown statically for 18-24 hours at 37°C. The following day, the overnight culture was centrifuged (4,000 x *g*, 10 min), and the bacterial pellet was washed once with PBS before being resuspended in 1 ml of complete DMEM (see Cell Culture section). The bacterial suspension was then normalized by optical density (OD_600_) to a concentration equivalent to 8×10⁸ CFU/ml (OD_600_ = 1) and further adjusted depending on the specific experimental application.

### Mouse wound excisional model

All *in vivo* procedures were approved by the Institutional Animal Care and Use Committee (IACUC) at Nanyang Technological University, Singapore (Protocol #ARF SBS/NIEA-0314), in accordance with national guidelines. Male C57BL/6J mice (6–7 weeks old, 22–25 g; InVivos, Singapore) were housed under specific-pathogen-free conditions. The wound infection model was adapted from a previous study^3^. Briefly, mice were anesthetized with 3% isoflurane, and dorsal hair was removed using clippers and a depilatory cream (Nair). The skin was disinfected with 70% ethanol, and a 6 mm full-thickness excisional wound was created using a sterile biopsy punch (Integra Miltex #33-36). The wound was immediately inoculated with 10 µl of an *E. faecalis* OG1RF suspension containing 2×10⁶ CFU. The wound was then sealed with a transparent dressing (Tegaderm, 3M #7100252702) to prevent contamination. Post-procedure, mice were housed individually to prevent wound disruption. At the experimental endpoint, mice were euthanized, and a 1 cm^2^ piece of skin tissue centered on the wound was excised. For bacterial enumeration, tissues were collected in sterile PBS, homogenized, and plated on BHI agar supplemented with rifampin to confirm infection by the inoculated strain. For molecular analysis, tissues were homogenized in either TRIzol reagent (Thermo Fisher #15596026) for RNA extraction or ice-cold RIPA buffer (Thermo Scientific #89900) supplemented with a protease inhibitor cocktail (Roche #11697498001) for protein extraction.

### scRNA-seq integration and downstream analysis

Single-cell datasets (GSE229257) were re-processed in R 4.3.2^63^ following the workflow established in our original study^22^. Raw matrices were imported with Seurat 5.1.0^64–67^. Cells expressing < 200 genes, > 6,000 genes or > 12 % mitochondrial reads were removed; genes detected in < 5 cells were discarded. Library size variation was normalised with SCTransform (method = “glmGamPoi”, vst.flavour = “v2”). Batch effects between biological replicates (uninfected vs. *E. faecalis*-infected wounds) were corrected with Seurat’s reciprocal PCA integration (30 PCs). Principal components (*n* = 30) were used for FindNeighbors/FindClusters (resolution = 0.4) and RunUMAP (dims = 1–30). Broad cell classes were assigned on canonical markers as previously reported^22^, sub-clustering of keratinocytes and fibroblasts employed a second round of PCA/UMAP at resolution = 0.6. All UMAPs use colour-blind-safe palettes generated with RColorBrewer 1.1-3^68^ (brewer.pal, palette = “Set2”).

Differential expression between infected and uninfected cells within each cluster was ranked by the Wilcoxon area-under-curve statistic using wilcoxauc^69^ (presto 1.0.0). Ranked lists served as input for gene-set enrichment analysis with fgseaMultilevel (fgsea 1.28.0, minSize = 15, maxSize = 5,000, eps = 0)^70,71^ Gene sets for unfolded protein response (UPR), oxidative-stress response (OSR) and heat shock response (HSR) were curated from MSigDB (v2023.1). Enrichment was considered significant at Benjamini–Hochberg FDR < 0.05^72^. For each cluster, the positive, significant normalised enrichment scores (NES) for infected cells were projected onto the UMAPs.

### RNA extraction, reverse transcription, and quantitative real-time PCR

Total RNA was extracted from cell lines and homogenised mice wound samples using the EZ-10 DNAaway RNA Miniprep Kit (Biobasic #BS88136) following the manufacturer’s protocol. RNA concentrations were quantified using Qubit Broad Range RNA Quantification Assay (Thermo Fisher #Q10210) together with a Qubit 3 Fluorometer following manufacturer’s protocol. Complementary DNA (cDNA) was synthesised from extracted RNA normalised to 1 µg per sample using RevertAid Reverse Transcriptase (Thermo Fisher #EP0441) following manufacturer’s protocol. qPCR was performed using Luna Universal qPCR Master Mix (New England Biolabs #M3003E) together with a CFX-96 (Bio-Rad) or a QuantStudio3 (Thermo Fisher) Real-Time PCR system following manufacturer’s protocol. Each 20 µl reaction has a final concentration of 2.5 ng/µl of cDNA and 0.25 µM of primer pairs (**Table S3**) for target genes. Relative mRNA was normalized to the housekeeping gene *Gapdh*/*GAPDH* using the 2^-ΔΔCt^ method^73^.

### Cell culture

Murine embryonic fibroblasts (NIH-3T3) and human keratinocytes (HaCaT) were cultured in DMEM (Gibco #11995065) supplemented with 10% heat-inactivated foetal bovine serum (FBS, Cytiva #SV30160.03) and 4 mM GlutaMAX (Gibco #35050061). This medium is referred to as "complete DMEM. The lentivirus packaging line, 293FT, was cultured in complete DMEM further supplemented with 0.1 mM MEM Non-Essential Amino Acids (Gibco #11140050), 6 mM L-glutamine (Gibco #25030081), 1 mM Sodium Pyruvate (Cytiva #SH30239.01), and 500 μg/ml Geneticin (Gibco Gibco #10131027). All cells were maintained at 37°C in a humidified 5% CO_2_ incubator. Cells were washed once with PBS (Gibco #14190144) and detached using 0.25% Trypsin-EDTA. Trypsinization times were 15 min for HaCaT, 5 min for 293FT, 4 min for 3T3R, and 3 min for NIH-3T3 cells. The reaction was neutralized with an equal volume of complete DMEM. Cells were pelleted by centrifugation, resuspended in fresh medium, and counted using a Countess 3 Automated Cell Counter. For experiments, cells were seeded in 12-well plates at densities of 2.85×10^4^ cells/cm^2^ (NIH-3T3), 1.15×10^4^ cells/cm^2^ (3T3R) or 1.15×10^5^ cells/cm^2^ (HaCaT) and incubated for 24 hours prior to use. For the transposon scree, 3T3R cells were seeded in 96-well plates at 2.85×10^4^ cells/cm^2^. Unless otherwise specified, the following final concentrations of reagents were used: tunicamycin at 0.2 µg/ml (for qPCR), 2.5 µg/ml (for scratch wound assays), or 5 µg/ml (for immunoblot/microscopy); the IRE1 inhibitor 4µ8c at 50 µM (MedChemExpress #HY-19707), added 1 hour prior to infection; H_2_O_2_ at 250 µM; catalase at 100 units/ml; and superoxide dismutase (SOD, Sigma-Aldrich #S5395) at 100 units/ml.

### Optimisation of *in vitro* infection

Assay conditions were optimised across (i) multiple timepoints and (ii) multiplicities of infection (MOI), to identify those that maximise *in vitro XBP1s* expression (data not shown).

### *In vitro* infection

Confluent NIH-3T3 and HaCaT cells were infected with *E. faecalis* at a MOI of 800 (800 CFU per host cell) and 600 (600 CFU per host cell), respectively. Following a 3-hour infection period, the medium was removed, and cells were washed three times with PBS. To eliminate extracellular bacteria, fresh complete medium supplemented with a gentamicin-penicillin antibiotic cocktail (50 µg/ml) was added, and the cells were incubated for an additional 21 hours^74^.

### Immunoblotting

Cells were lysed with RIPA buffer (Thermo Fisher #89901) supplemented with reconstituted cOmplete protease inhibitor cocktail (Roche #11697498001) by gentle agitation on ice for 5 min before centrifugation for 15 min at 12,000 x *g* at 4°C. A mixture of 15 µg of total proteins was separated on 10% SDS-PAGE and transferred on nitrocellulose membranes. Immunoblotting was performed with appropriate primary antibodies and IRDye-conjugated secondary antibodies (**Table S4**). Proteins were visualized using the NIR fluorescence system (Odyssey CLx Imaging System).

### *In vitro* scratch wound assay model

Scratch assays were performed in 12-well plates, adapting a previously published protocol to facilitate automated microscopy^75^. Confluent HaCaT cell monolayers were scratched with a sterile P200 pipette tip and subsequently infected as described in the *In vitro* infection section. Following the post-infection wash, fresh complete medium supplemented with antibiotics and 25 mM HEPES (Gibco #15630080) was added to each well. Wound closure was monitored on a Zeiss Axio Observer 7 microscope (10x magnification), acquiring brightfield images every 30 min for 45 h in a controlled environment (37°C, 5% CO_2_). The resulting time-lapse images were analyzed using a customised CellProfiler pipeline to quantify the scratch area^76^. To ensure accuracy, images where the wound area was misidentified by the automated pipeline were manually measured in ImageJ using the Wound Healing Size Tool plugin^77^.

### XBP1s reporter cell line

pLVX-XBP1-mNeonGreen-NLS plasmid was a gift from David Andrews (**Table S5**). Codon-optimized mApple cDNA was synthesized as a gBLOCK fragment (IDT) and inserted in pLVX-XBP1-mNeonGreen-NLS to replace mNeonGreen using Gibson Assembly (New England Biolabs #E2611L) according to manufacturer protocol. The resulting plasmid, pLVX-XBP1-mApple-NLS, was sequence-verified and then co-transfected into 293FT cells with the pLP1, pLP2, and pLP/VSVG packaging plasmids (**Table S5**). Supernatants containing lentiviral particles were harvested at 36- and 60-hour post-transfection, pooled, and filtered (0.45 μm). NIH-3T3 cells were then transduced with the filtered virus for 24 hours in the presence of 8 µg/ml of polybrene (Sigma-Aldrich #H9268). After 24 hours recovery, infected cells were selected with 2 µM puromycin. Clonal cell lines were established by seeding single cells into 96-well plates and expanding them for two weeks. Finally, positive clones were validated by assessing homogenous fluorescent signal upon tunicamycin treatment. One validated clone, designated the 3T3 reporter (3T3R) line, was selected for this study.

### Transposon screen

A high-throughput screen was performed using an established *E. faecalis* OG1RF mariner transposon library containing 14,976 mutants arrayed in 96-well plates^78^. Following overnight growth, the optical density (OD_600_) of each mutant culture was measured with a Tecan M200 microplate reader. For infection, 5 µl of each culture was added to 3T3R cells seeded in 96-well plates, and plates were centrifuged at 300 x *g* for 5 minutes to synchronize contact. Following infection, cells were stained with 2.5 µg/ml Hoechest 33342 (Thermo fisher #H21492) for 15 minutes, and the medium was then replaced with phenol-red free DMEM (Gibco #31053028) supplemented with 25 mM HEPES, 1 mM sodium pyruvate, and 4 mM GlutaMAX. Plates were imaged on a Zeiss CellDiscoverer 7 microscope (10x magnification) using two fluorescence channels to detect the XBP1s-mApple reporter (Ex/Em: 570/594 nm) and Hoechst-stained nuclei (Ex/Em: 348/455 nm). The images were subsequently analysed with CellProfiler 4.2.1 to quantify the percentage of UPR-positive (mApple-expressing) cells in each well.

### Construction of in-frame deletion mutants in *E. faecalis*

General molecular biology reagents were sourced as follows: genomic DNA from *E. faecalis* was isolated using the Wizard Genomic DNA Purification Kit (Promega #A1120), while plasmid DNA was isolated from *E. coli* using the PureLink Plasmid Miniprep Kit (Invitrogen #K210011). All primers used in this study (**Table S6**) were designed based on the *E. faecalis* OG1RF genome (NC_017316). Gene fragments were amplified with Phusion High-Fidelity DNA Polymerase (Thermo Scientific #F530), and routine screening was performed with Taq DNA Polymerase (NEB #M0273). T4 DNA ligase and all restriction enzymes used according to the manufacturer’s protocols (NEB). In-frame deletion mutants were generated using the temperature-sensitive shuttle vector pGCP213 (**Table S5**), following a previously described protocol^79^. Deletion constructs were created using two main strategies. For most single genes and smaller operons, regions of approximately 450 bp flanking the target were amplified from OG1RF gDNA; the upstream region was amplified with primer pair P1/P2 and the downstream region with P3/P4. These fragments were then fused by overlap extension PCR using the outer primers P1 and P4 and subsequently cloned into the *PstI* site of pGCP213. For the larger ΔEET and ΔT7SS operons, a gBlock Gene Fragment (IDT) containing the fused upstream and downstream flanking regions was synthesized and cloned into the vector. The resulting deletion constructs were transformed into the appropriate *E. faecalis* parent strain by electroporation. Transformants were first selected on BHI-erythromycin agar at the permissive temperature of 30°C. To promote chromosomal integration, colonies were then passaged at the non-permissive temperature of 42°C with erythromycin selection. Curing of the integrated plasmid was achieved by passaging the bacteria at 30°C in antibiotic-free BHI. Finally, the successful deletion of the target gene or operon was verified by colony PCR using external (Screen F/R) and internal (Intern F/R or Intern R) primer pairs (**Table S6**).

### XBP1s fluorescent reporter assay

Following infection or treatment, 3T3R cells were stained with Hoechst 33342 (2.5 µg/ml) and transferred to a phenol-red-free imaging medium. For each well, a 3×3 grid of images was acquired on a Zeiss CellDiscoverer 7 microscope. The intensity of the XBP1s-mApple signal within each nucleus (identified by Hoechst staining) was quantified using a CellProfiler pipeline, and the average intensity per well was used to gauge the level of UPR induction.

### Growth curve assay

To assess bacterial growth, overnight bacterial cultures were processed as described in bacterial strains and growth conditions. After pelleting, they were normalised to a starting OD_600_ of 0.05 in phenol red free complete media on a 96-well plate which was sealed with a Breathe-Easy sealing membrane (Sigma-Aldrich #Z380059) following manufacturer’s protocols. OD_600_ readings were taken at 10-min intervals over 20 h. The rate of change of the OD_600_ readings was calculated at 50-min intervals and the highest rate of change was used to calculate the doubling time.

### Antibiotic time-kill assay

To assess antibiotic killing, the supernatant of infected HaCaT cells were collected at 4, 5, 6, and 24 hpi. These were serially diluted on 96-well plates and 5 µl of the dilutions were spotted onto BHI-Agar plates which were incubated at 37°C for 24 hours. Plates were imaged using a ProtoCOL3 Plus system (Don Whitely Scientific) and bacterial colonies were manually enumerated on ImageJ using the multi-point tool.

### Cytotoxicity assay

To quantify total cytotoxicity, both detached (floating in the medium) and attached cells were collected and analyzed from infected HaCaT cell cultures. First, to collect the detached cell fraction, the culture medium was harvested, and each well was washed once with 1 ml of PBS. This wash was pooled with the collected medium, and the mixture was centrifuged (300 x *g*, 5 min). The resulting cell pellet was carefully resuspended in 20 µl of complete medium. Next, to collect the attached cell fraction, the remaining cells in the well were trypsinised for 15 min, neutralized with complete medium, pelleted by centrifugation, and resuspended in 500 µl of fresh complete medium. The viability of both the detached and attached cell suspensions was determined separately using a Countess 3 Automated Cell Counter with trypan blue staining. Total cytotoxicity was then calculated by summing the number of non-viable cells from both fractions and dividing by the total number of cells (viable and non-viable) from both fractions (**Equation 1**).

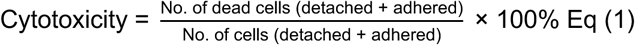

### Superoxide assay

Extracellular superoxide generation was measured by adapting a previously published cytochrome C reduction assay for a 96-well plate format^18^. Briefly, *E. faecalis* cultures were normalized to an an OD_600_ of 0.005 in 200 µl of phenol red free complete media containing 20 µM of cytochrome C (Sigma-Aldrich #C3131). The reduction of cytochrome C was measured as the change in absorbance at 550 nm (with 650 nm as the reference wavelength) every 2 minutes for 90 minutes at 37°C using a Tecan M200 microplate reader. The rate of superoxide generation was calculated from the maximal rate of change in absorbance, after correcting for pathlength. This rate was determined using Beer’s Law with an extinction coefficient of 21.5 mM⁻¹cm⁻¹ for reduced cytochrome C (**Equation 2**). To determine the amount of superoxide specifically, the rate measured in a parallel reaction containing 100 units/ml of superoxide dismutase (SOD; Sigma-Aldrich #S5395) was subtracted from the rate measured in its absence.

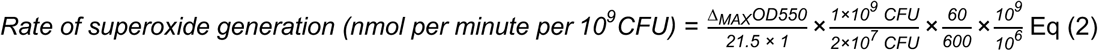

### Lipid peroxidation assay

Lipid peroxidation was assessed in HaCaT cells using the Image-iT Lipid Peroxidation Kit (Thermo Fisher #C10445) at 24 hpi following manufacturer’s protocols. Imaging was performed on a Zeiss CellDiscoverer 7 microscope using two fluorescence channels to detect fluorescence from reduced (Ex/Em: 592/614 nm) and oxidised (Ex/Em: 495/519 nm) BODIPY 581/591 C11. The mean intensity of the reduced and oxidised fluorescent reporters was quantified using a CellProfiler pipeline, with lipid peroxidation calculated based on the ratio of reduced:oxidised signals for each condition.

### Statistics

Statistical analyses were performed using GraphPad Prism 9 and 10. In bar-dot plots, dots represent individual replicates, and the bar height indicates the median. Statistical significance was determined using either a one-way ANOVA with Dunnett’s multiple comparisons test or, for scratch wound assays, a two-way ANOVA with Tukey’s multiple comparisons test. An adjusted *p*-value < 0.05 was considered significant. Unless otherwise stated in the figure legends, all experiments were performed with a minimum of three independent replicates.

## SUPPLEMENTAL INFORMATION

**Document S1. Figures S1-S4, Tables S2-S6**

**Table S1. Stress Response Gene Set.** Related to Figure 1. Excel Spreadsheet.

**Video S1. *E. faecalis* infection and tunicamycin treatment impair HaCaT cell migration**. Related to Figure 2E.

**Video S2. IRE1 inhibition alters HaCaT cell migration in uninfected and *E. faecalis*-infected conditions.** Related to Figure 2G.

**Video S3. The ΔEET mutant does not impair HaCaT cell migration.** Related to Figure 3E.

**Video S4. The effect of IRE1 inhibition on HaCaT cell migration during infection with WT or ΔEET *E. faecalis*.** Related to Figure 3F.

**Video S5. Catalase rescues H_2_O_2_-induced migration defects in uninfected HaCaT cells.** Related to Figure 4K.

**Video S6. Catalase restores migration in *E. faecalis*-infected HaCaT cells.** Related to Figure 4L.

