## Supplementary material for "*Enterococcus faecalis* redox metabolism activates the unfolded protein response to impair wound healing": Document S1

### SUPPLEMENTAL FIGURES

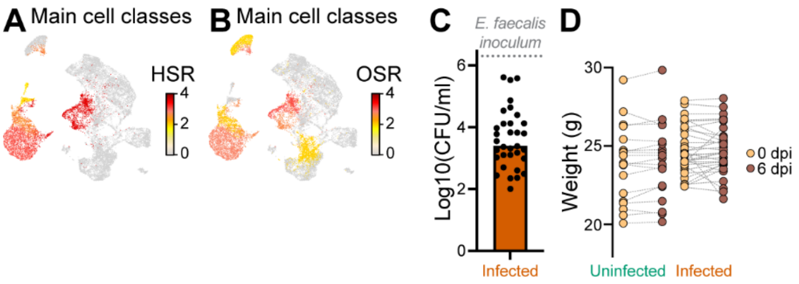


**Figure S1. Stress response analysis and characterization of the *E. faecalis* wound infection model. Related to Figure 1.**
**(A**) Per-cell enrichment score for a curated heat shock response (HSR) gene set projected onto the uniform manifold approximation and projection (UMAP) in Fig. 1A. **(B**) Per-cell enrichment score for a curated oxidative stress response (OSR) gene set projected onto the UMAP in Fig. 1A. **(C)** Mouse wounds were inoculated with 2x10$^{6}$ colony forming units (CFU) of *E. faecalis* WT. The CFU were quantified at 6 days post-infection (dpi) to quantify *E. faecalis* burden at the wound site (*n* = 33). **(D)** Weight of uninfected and WT-infected 6–7-week-old C57BL/6J mouse skin wounds from the day of wounding (0 dpi) until 6 dpi (uninfected, *n* = 20; WT, *n* = 33).

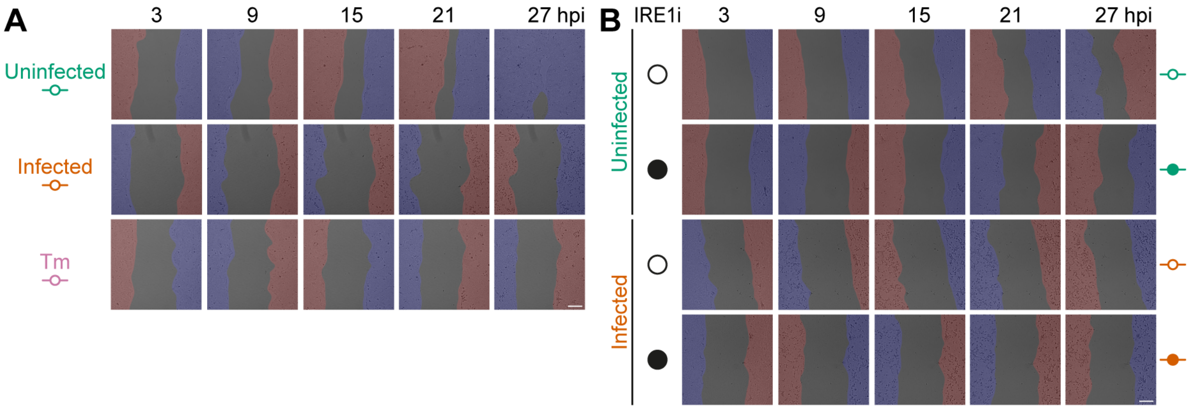


**Figure S2. *In vitro* epithelial cell migration following infection and UPR modulation. Related to Figure 2.**
**(A-B)** Representative scratch wound images for (A) uninfected, infected and Tm-treated cells, which were also (B) treated with 0.5% DMSO control (open circles) or IRE1i (close circles). Scale bars, 200 µm. Areas highlighted in red and blue demarcate the cell monolayer identified by Cellprofiler.

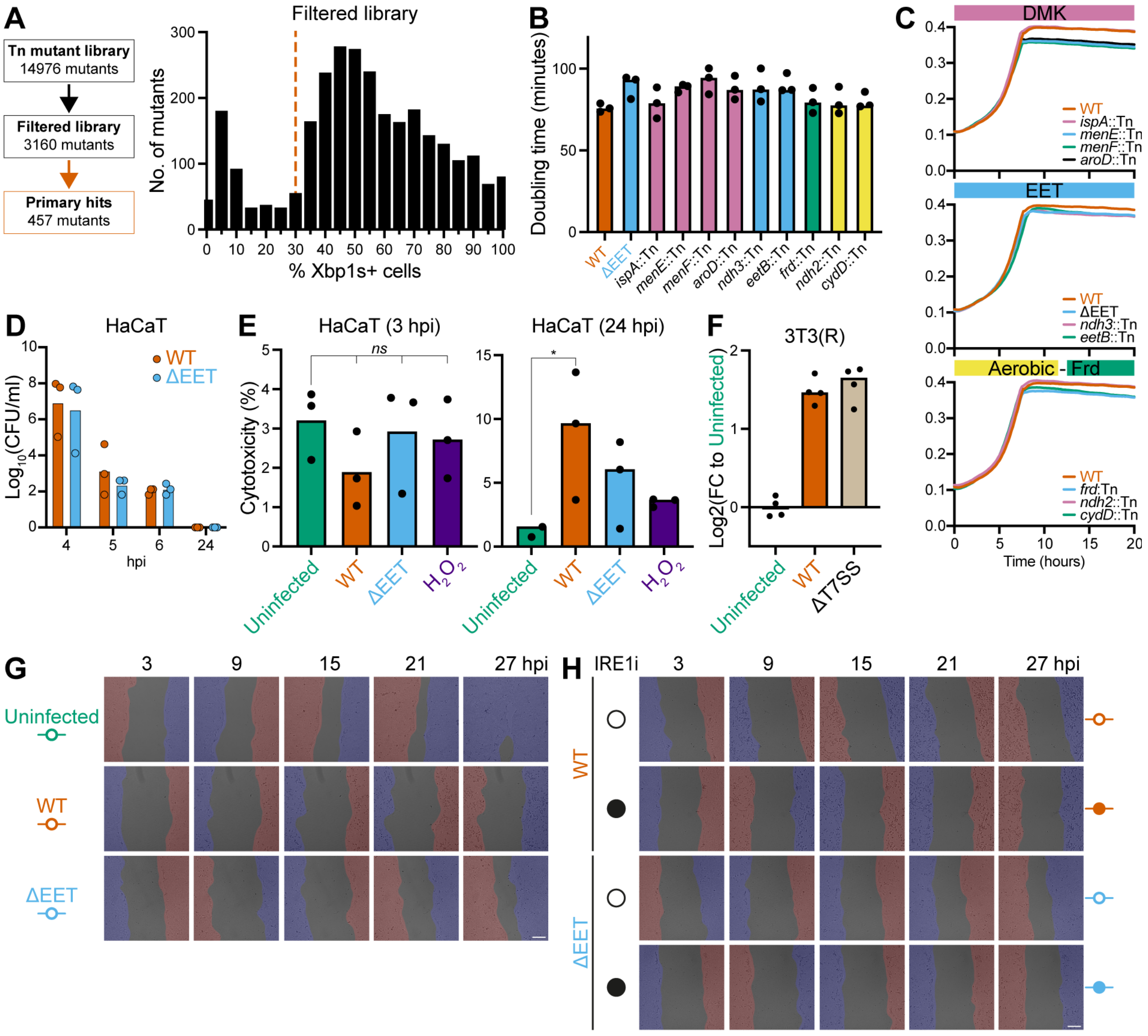


**Figure S3. Selection and characterization of E. faecalis mutants that fail to induce the host UPR. Related to Figure 3.**

**(A)** Filtering and thresholding (orange dotted line) the transposon mutant library to identify primary hits for further validation. Mutants were filtered using the following criteria: (i) Contains only one transposon insertion in coding sequence, (ii) > 10 H3342+ cells in an image, (iii) > 0 Xbp1s+ cells in an image (iv) Overnight culture with an absorbance $OD_{600}$ > 0.3. **(B-C)** Doubling times (B) and growth curves (C) of WT and UPR-defective mutants, determined by monitoring absorbance ($OD_{600}$) during growth in cell culture medium at 37°C. **(D)** Survival of extracellular WT and ΔEET during antibiotic exposure in HaCaT co-cultures. **(E)** Cytotoxicity at 3 and 24 hpi after exposing HaCaT to WT, ΔEET, or 250 μM $H_2O_2$ using a modified trypan blue assay. **(F)** UPR induction by WT and ΔT7SS in a 3T3(R) background. **(G-H)** Representative scratch wound images for (G) uninfected, WT-infected and ΔEET-infected cells, which were also (H) treated with 0.5% DMSO control (open circles) or IRE1i (close circles). Significance was determined using one-way ANOVA Dunnett's test (E) (*ns*, non-significant; **p* < 0.05). Scale bars; 200 μm. Areas highlighted in red and blue demarcate the cell monolayer identified by Cellprofiler.

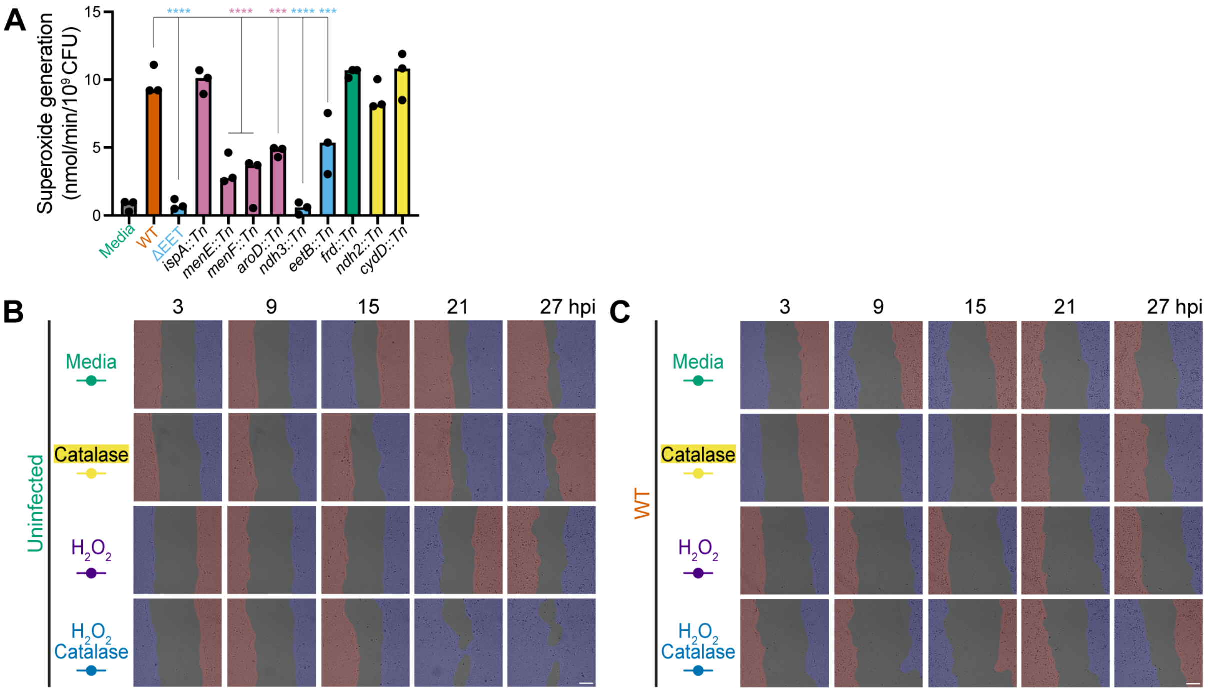


**Figure S4. Representative scratch wound images in uninfected and WT-infected conditions, when treated with catalase and/or hydrogen peroxide. Related to Figure 4.**

**(A)** Superoxide generation rate of UPR defective mutants. Significance was determined using one-way ANOVA Dunnett's test ($n$ = 3, ***$p$ < 0 .001, ****$p$ < 0.0001). **(B-C)** Representative scratch wound images in uninfected (B) or WT-infected (C) HaCaT cells treated with catalase and/or $H_2O_2$. Scale bars, 200 µm. Areas highlighted in red and blue demarcate the cell monolayer identified by Cellprofiler.

**SUPPLEMENTAL TABLES**

**Table S1. Stress Response Gene Set.** Related to Figure 1. Excel Spreadsheet.

**Table S2. Bacterial strains.** Related to Figure 1-4.

| Strain | Description | Reference or source |
|---|---|---|
| ***E. faecalis*** | | |
| OG1RF | Laboratory strain, $Rif^R$, $Fus^R$ | Dunny et al.[1] |
| ΔEET | EET operon chromosomal deletion mutant, $Rif^R$, $Fus^R$ | This study |
| Δ*ndh3* | *ndh3* chromosomal deletion mutant, $Rif^R$, $Fus^R$ | This study |
| Δ*eetB* | *eetB* chromosomal deletion mutant, $Rif^R$, $Fus^R$ | This study |
| ΔMEN | MEN operon chromosomal deletion mutant, $Rif^R$, $Fus^R$ | This study |
| Δ*menE* | *menE* chromosomal deletion mutant, $Rif^R$, $Fus^R$ | This study |
| Δ*menF* | *menF* chromosomal deletion mutant, $Rif^R$, $Fus^R$ | This study |
| ΔT7SS | T7SS operon chromosomal deletion mutant, $Rif^R$, $Fus^R$, $Cm^R$ | This study |
| *ispA*::*tn* | *ispA* transposon insertion mutant (Tn insertion at 740028), $Rif^R$, $Fus^R$, $Cm^R$ | Kristich *et al.* (2008)[2] |
| *menE*::*tn* | *menE* transposon insertion mutant (Tn insertion at 344801), $Rif^R$, $Fus^R$, $Cm^R$ | Kristich *et al.* (2008)[2] |
| *menF*::*tn* | *menF* transposon insertion mutant (Tn insertion at 345749), $Rif^R$, $Fus^R$, $Cm^R$ | Kristich *et al.* (2008)[2] |
| *aroD*::*tn* | *aroD* transposon insertion mutant (Tn insertion at 1501457), $Rif^R$, $Fus^R$, $Cm^R$ | Kristich *et al.* (2008)[2] |
| ndh3::*tn* | *ndh3* transposon insertion mutant (Tn insertion at 2658965), $Rif^R$, $Fus^R$, $Cm^R$ | Kristich *et al.* (2008)[2] |
| *eetB*::*tn* | *eetB* transposon insertion mutant (Tn insertion at 2660397), $Rif^R$, $Fus^R$, $Cm^R$ | Kristich *et al.* (2008)[2] |
| *frd*::*tn* | *frd* transposon insertion mutant (Tn insertion at 2043345), $Rif^R$, $Fus^R$, $Cm^R$ | Kristich *et al.* (2008)[2] |
| *ndh2*::*tn* | *ndh2* transposon insertion mutant (Tn insertion at 1729431), $Rif^R$, $Fus^R$, $Cm^R$ | Kristich *et al.* (2008)[2] |
| *cydD*::*tn* | *cydD* transposon insertion mutant (Tn insertion at 1735842), $Rif^R$, $Fus^R$, $Cm^R$ | Kristich *et al.* (2008)[2] |
| ***E. coli*** | | |
| Stellar | *E. coli* host strain for routine cloning | Laboratory stock |
| DH5α | *E. coli* host strain for routine cloning | Laboratory stock |

**Table S3. Oligonucleotide primers used in this study.** Related to Figure 1-4.

| Species | *Gene* | Forward (5' to 3') | Reverse (5' to 3') |
|---|---|---|---|
| Human | *CHOP* | GGA AAC AGA GTG GTC ATT CCC | CTG CTT GAG CCG TTC ATT CTC |
| | *EDEM1* | ACG AGC AGT GAA AGC CCT TTG G | CCA CTC TGC TTT CCA ACC CAG T |
| | *GAPDH* | GCC ATC AAT GAC CCC TTC ATT | TCT CGC TCC TGG AAG ATG G |
| | *HERPUD1* | TCG TGG TTC TAA TCG GGG ACA | CCA GGG GAA GAA AGG TTC CG |
| | *XBP1s* | TGC TGA GTC CGC AGC AGG TG | GCT GGC AGG CTC TGG GGA AG |
| Mouse | *Chop* | AAG ATG AGC GGG TGG CAG CG | GCT CCC AGC TGG ACA CCG TC |
| | *Gapdh* | TCA GGA GAG TGT TTC CTC GTC CC | TCT CGG CCT TGA CTG TGC CG |
| | *Herpud1* | TTG GAG CTG AGT GGC GAC CG | GGA AGC AAA TCT TGG AGA CAC TGG T |
| Omni[3] | *XBP1s* | GCT GAG TCC GCA GCA GGT | CWG GGT CCA ACT TGW MCA GAA T |
| | *GAPDH* | ACC ATC TTC CAG GAG CGA GA | GGG CCA TCC ACA GTC TTC TG |

**Table S4. Antibodies used in this study.** Related to Figure 2.

| Target | Host | Dilution for IB | Reference or source |
|---|---|---|---|
| XBP1s/XBP1u | Rabbit | 1:500 | Cell signalling technology #12782 |
| BiP | Rabbit | 1:1,000 | Cell signalling technology #3177 |
| α-TUBULIN | Mouse | 1:10,000 | DSHB #12G10 |
| IRDye 800CW Goat anti-Rabbit IgG | Goat | 1:15,000 | Licor #926-32211 |
| IRDye 680LT Goat anti-Mouse IgG | Goat | 1:15,000 | Licor # 926-68020 |

**Table S5. Plasmids used in this study.** Related to Figure 3-4.

| Plasmid | Relevant characteristics | Reference or source |
|---|---|---|
| pLVX-Xbp1s-mApple | Lentiviral transfer plasmid for XBP1 splicing reporter | This study |
| pLP1 | Lentiviral packaging plasmid | Thermo Fisher #K497500 |
| pLP2 | Lentiviral packaging plasmid | Thermo Fisher #K497500 |
| pLP/VSVG | Lentiviral packaging plasmid | Thermo Fisher #K497500 |
| pCGP213 | Thermosensitive shuttle plasmid | Nielsen *et al.* (2012)[4] |

**Table S6. Primers used to generate and validate clean deletion *E. faecalis* mutants.** Related to Figure 3-4.

| Mutant | Primers | 5' to 3' |
|---|---|---|
| ΔEET | Oligo P1 | GCG TAA TCA TAT CGA TGG TCA TAG C |
| | Oligo P2 | CTG GAA TTC TGC AGA TAT CCA TCA C |
| | Oligo P3 | TCT GCA GAA TTC CAG GCC AAG CAA CGC ACT AAT |
| | Oligo P4 | TCG ATA TGA TTA CGC ATT TTC TCT TGT CAA AAT CGT TTG T |
| | Screen F | CGA ACG AAC CAG AAC CAG CC |
| | Screen R | AAC GGG TCA GCA TTA TCG GG |
| | Intern F | GCG GGT GTT CTC GGA TTT TGG |
| | Intern R | CTT AAC GGT TCA AAC CGG CTG G |
| Δ*ndh3* | Oligo P1 | GAT TGT CAG CAA TCG ATA ATC GAT TGA CA |
| | Oligo P2 | GGC GGC CGT TAC TAG TGC GAC TTT TCT TAA TGA ATT CTT TGA |
| | Oligo P3 | TAC CGA GCT CGG ATC CTC ACA AAC ATC AAA AGT TTG TGA TAC A |
| | Oligo P4 | CGA TTG CTG ACA ATC CGC TCC TTA ATA TTG T |
| | Screen F | AGC AAT CGA TAA TCG ATT GAC A |
| | Screen R | GCG ACT TTT CTT AAT GAA TTC TTT GA |
| Δ*eetB* | Oligo P1 | GTC ATG GAC TGC AGC TTG AA |
| | Oligo P2 | ATT GCT ATG CTC GTT GAA CGA AAC ATA AGT GGA G |
| | Oligo P3 | CTC CAC TTA TGT TTC GTT CAA CGA GCA TAG CAA TAT A |
| | Oligo P4 | GGT GAT CGG ATC CGC GTG AA |
| | Screen F | GGT ATT CCA CAT AGG ATG TA |
| | Screen R | GTT TAT CAC CGT GGA AGC T |
| ΔMEN | Oligo P1 | CAC CTA AAC TGC AGG CAG TTT C |
| | Oligo P2 | GAC TGT ATT TTA GCA AAT TCA TTC TTT AAT TGT TGT CCA G |
| | Oligo P3 | CTG GAC AAC AAT TAA AGA ATG AAT TTG CTA AAA TAC AGT C |
| | Oligo P4 | CGG GAT CCC GCT ACC ACT CAT TTT AA |
| | Screen F | CAG TCC TTT CTA ATA AAA GAG AG |
| | Screen R | GTC CTA ATT GTA AAT GGT GAA |
| | Intern R | GCA CCA ATA ATC GTT TTA AC |
| Δ*menE* | Oligo P1 | CAC GTT TAA ATG TTC TAG ATT TAC AAC G |
| | Oligo P2 | GCC TCT CTT TCT TCT GTC TTG TAT AGT AAA GAA GTG |
| | Oligo P3 | CACT TCT TTA CTA TAC AAG ACA GAA GAA AGA GAG GCG T |
| | Oligo P4 | GAG CCC CGG TAC CAG AAA TAG |
| | Screen F | GAT GTG ACA ATG GAG TGG GC |
| | Screen R | CCT TGG TAT GCC TTT TCA CC |
| | Intern R | CTT TAA AGC TGG CGC TGT AG |
| Δ*menF* | Oligo P1 | CAT GGG CTA TCT AGA TGC ACA AG |
| | Oligo P2 | CCT CGT AAC ATC GGT AGG CTT GTC GTA ATT C |
| | Oligo P3 | GAA TTA CGA CAA GCC TAC CGA TGT TAC GAG GAA TTG |
| | Oligo P4 | GTT TGC AGC TGC AGT TCC AGA G |
| | Screen F | CAA GAC CAG TGT TCA CGA AA |
| | Screen R | GAA CGT TCA TCG ACA TCG AC |
| | Intern R | CCT TGT CAA TCT TTG TTT CCG |
| ΔT7SS | Oligo P1 | GCG TAA TCA TAT CGA TGG TCA TAG C |
| | Oligo P2 | CTG GAA TTC TGC AGA TAT CCA TCA C |
| | Oligo P3 | TCT GCA GAA TTC CAG AAA ATT TTA TCA ATT GGC AAT CA |
| | Oligo P4 | TCG ATA TGA TTA CGC CCA ATT TTC GGT GTT CAC AGC CTG |
| | Screen F | GGG AAT GGC ACC CTG AAA GA |
| | Screen R | CTT CGC GCT TGG CTT TTT GA |
